# Recruitment of CRISPR-Cas systems by Tn7-like transposons

**DOI:** 10.1101/154229

**Authors:** Joseph E. Peters, Kira S. Makarova, Sergey Shmakov, Eugene V. Koonin

## Abstract

A survey of bacterial and archaeal genomes shows that many Tn7-like transposons contain ‘minimal’ type I-F CRISPR-Cas systems that consist of fused cas8f and cas5f, cas7f and cas6f genes, and a short CRISPR array. Additionally, several small groups of Tn7-like transposons encompass similarly truncated type I-B CRISPR-Cas systems. This gene composition of the transposon-associated CRISPR-Cas systems implies that they are competent for pre-crRNA processing yielding mature crRNAs and target binding but not target cleavage that is required for interference. Here we present phylogenetic analysis demonstrating that evolution of the CRISPR-Cas containing transposons included a single, ancestral capture of a type I-F locus and two independent instances of type I-B loci capture. We further show that the transposon-associated CRISPR arrays contain spacers homologous to plasmid and temperate phage sequences, and in some cases, chromosomal sequences adjacent to the transposon. A hypothesis is proposed that the transposon-encoded CRISPR-Cas systems generate displacement (R-loops) in the cognate DNA sites, targeting the transposon to these sites and thus facilitating their spread via plasmids and phages. This scenario fits the “guns for hire” concept whereby mobile genetic elements can capture host defense systems and repurpose them for different stages in the life cycle of the element.

**Importance:** CRISPR-Cas is an adaptive immunity system that protects bacteria and archaea from mobile genetic elements. We present comparative genomic and phylogenetic analysis of degenerate CRISPR-Cas variants associated with distinct families of transposable elements and develop the hypothesis that such repurposed defense systems contribute to the transposable element propagation by facilitating transposition into specific sites. Such recruitment of defense systems by mobile elements supports the “guns for hire” concept under which the same enzymatic machineries can be alternately employed for transposon proliferation or host defense.

## Introduction

CRISPR (Clustered Regularly Interspaced Short Palindromic Repeat)-Cas (CRISPR-ASsociated proteins) systems are the only adaptive immune systems identified in prokaryotes (1). CRISPR-Cas systems possess modular organization which roughly reflects the three main functional stages of the CRISPR immune response: i) spacer acquisition (known as adaptation), ii) pre-crRNA processing and iii) interference (1). CRISPR-Cas systems are highly diverse but can be partitioned into two distinct classes based on the organization of the effector module that is responsible for processing and adaptation (2). Class 1 CRISPR-Cas systems are further divided into 3 types and 12 subtypes in all of which the effector modules are multisubunit complexes of Cas proteins (2). In contrast, in the currently identified 3 types and 12 subtypes of Class 2, the effector modules are represented by a single multidomain protein, such as the thoroughly characterized Cas9 (3-5).

At the adaptation stage, the Cas1-Cas2 protein complex, in some instances with additional involvement of effector module proteins, captures a segment of the target DNA (known as the protospacer) and inserts it at the 5’end of a CRISPR array (6-9). In the second, processing stage, a CRISPR array is transcribed into a long transcript known as pre-CRISPR (cr) RNA that is bound by Cas proteins and processed into mature, small crRNAs. In most Class 1 systems, the pre-crRNA processing is catalyzed by the Cas6 protein that, in some cases, is loosely associated with the effector complex (1, 10). The final, interference step involves binding of the mature crRNA by the multisubunit effector complex, scanning a DNA or RNA molecule for a sequence matching the crRNA guide and containing a protospacer adjacent motif (PAM), and cleavage of the target by a dedicated nuclease domain(s) (1, 10-12). The identity of this nuclease(s) differs between type I and type III CRISPR-Cas systems. In type I, the protein responsible for target cleavage is Cas3 which typically consists of a Superfamily II helicase and HD-family nuclease domains (13). After the effector complex, which is denoted Cascade (CRISPR-associated complex for antiviral defense (14)) in type I systems, recognizes the cognate protospacer in the target DNA, it recruits Cas3, after which the helicase unwinds the target DNA duplex, and the HD nuclease cleaves both strands (15, 16). Type III systems lack Cas3, and the protein responsible for target cleavage is Cas10 which contains a polymerase-cyclase and HD-nuclease domains that are both required for the target degradation (17, 18).

In some of the CRISPR-Cas systems, the adaptation genes are encoded separately or even are missing from the genome containing effector complex genes. Among these non-autonomous CRISPR-Cas systems, those of type III systems have been characterized in most detail (1). It has been shown that class III effector complexes can utilize crRNA originating from CRISPR arrays associated with type I systems and thus do not depend on their own adaptation modules (19-23). Furthermore, the CRISPR-Cas systems of type IV, that are often encoded on plasmids, typically consist of the effector genes only (2). No adaptation genes and no associated nuclease domains could be found in the type IV loci although, occasionally, CRISPR arrays and *cas6*-like genes are present. The type IV systems have not yet been studied experimentally, so their mode of action remains unknown. Finally, several variants of type I systems, similarly to type IV, lack adaptation genes and genes for proteins involved in DNA cleavage. A “minimal” variant of subtype I-F has been identified in the bacterium *Shewanella putrefaciens*, with an effector module that consists only of Cas5f, Cas6f and Cas7f proteins, and lacks the large and small subunit present in other Cascade complexes (24). Even more dramatic minimization of subtype I-F has been observed in another variant of subtype I-F that lacks the adaptation module and consists solely of three effector genes, namely a fusion of *cas8f* (large subunit) with *cas5f* that is unique for this variant, *cas7f* and *cas6f* (Figure 1A) (2). Given the composition of their Cascade complex, these Cas1-less minimal subtype I-F systems can be predicted to process pre-crRNA, yielding mature crRNAs, and recognize the target. However, they lack the Cas3 protein and therefore cannot be expected to be competent for target cleavage. Here we report an exhaustive *in silico* analysis of this system showing that it is strongly linked to a specialized group of transposons related to the well-studied Tn7.

**Figure 1.**
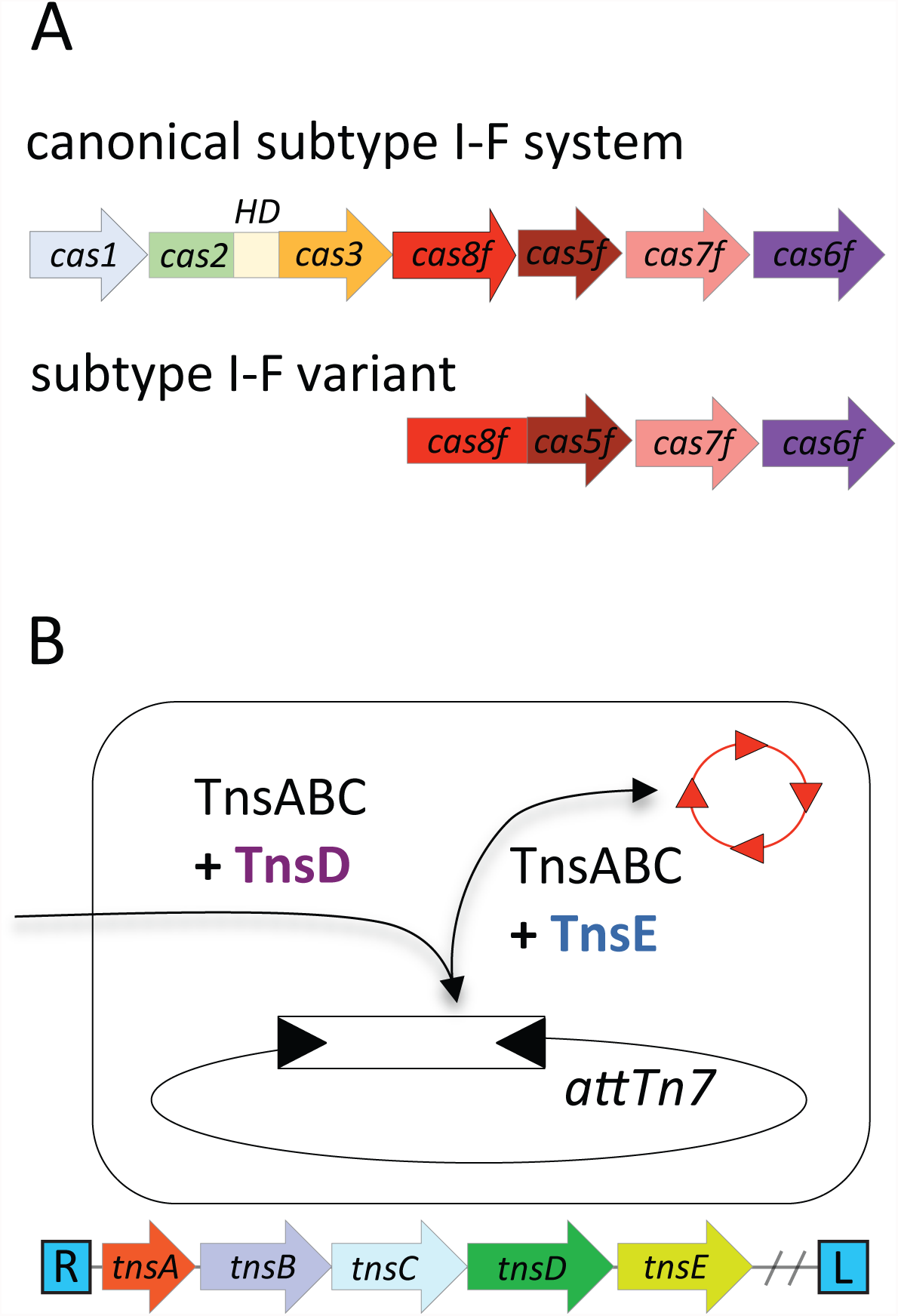
Schematic representation of the complete and minimal type I-F CRISPR-Cas systems and Tn7 transposition. A. Gene organizations of a complete and a minimal type I-F CRISPR-Cas system lacking the genes for proteins responsible for adaptation and target cleavage. Minimal I-F systems contain fused *cas8f* and *cas5f* genes that are characteristic of this group (2). Together, these proteins can be predicted to be subunits of a minimal Cascade complex. B. Gene structure of the Tn7 genes flanked by left (L) and right (R) end sequences. Transposition catalyzed by the TnsABC+TnsD proteins directs the transposon into a single chromosomal site in bacterial genomes (*attTn7*). Transposition catalyzed by the TnsABC+TnsE proteins preferentially directs transposition into actively conjugating DNA and filamentous bacteriophage (shown by a red circle with arrows). The transposon is denoted by a rectangle in the attachment site. DNA sequence omitted in the graphic is denoted with two back slashes. See text for details.

The canonical Tn7 is notable because of the level of control it exerts over the target site selection (25). Three transposon-encoded proteins form the core transposition system including a heteromeric transposase (TnsA and TnsB) and a regulator protein (TnsC) (Figure 1B). In addition to the core TnsABC transposition proteins, Tn7 elements encode dedicated target site selection proteins (TnsD and TnsE) that only allow transposition when specific types of target sites are available. In conjunction with TnsABC, the sequence-specific DNA-binding protein TnsD directs transposition into a conserved site referred to as the Tn7 attachment site, *attTn7* (26). Transposition mediated by TnsABC + TnsE is preferentially directed into mobile plasmids and bacteriophages owing to the ability of TnsE to recognize complexes formed during specific types of DNA replication (27-30). Mobile elements related to Tn7 and encoding proteins homologous to TnsA, TnsB, and TnsC have been described that appear to use distinct attachment sites recognized by TnsD/TniQ-like proteins, but do not encode a TnsE-like protein (31).

Our analysis shows that the Cas1-less I-F systems associate with a distinct group of Tn7-like elements. These transposons encode TnsD/TniQ-like proteins and utilize novel attachment sites but lack TnsE-like proteins that normally promote horizontal transfer of the elements. Several identified matches for the spacers from the transposon-associated CRISPR arrays suggest that this system might function by targeting transposition to target sites enabled by guide crRNAs. We hypothesize that the subtype I-F CRISPR-Cas machinery recruited by these elements facilitates their horizontal dissemination, mostly via plasmids and/or phages.

## Results and discussion

### A variant of the type I-F CRISPR-Cas system is specifically associated with a distinct family of Tn7-like elements

For the purpose of comprehensive identification type I-F CRISPR-Cas loci, we chose the Cas7f protein as the probe given that it is the most conserved component in all systems of this subtype including the “minimal” variant lacking *cas1* and *cas2-cas3* genes. Using a PSI-BLAST search started with Cas7f profiles, we obtained 2905 Cas7f protein sequences, mapped them onto the respective genomes and annotated the genes in the 10 kb up- and downstream neighborhoods of the *cas7f* genes using PSI-BLAST against the conserved domain database (CDD). These 20 kb loci are long enough to cover a typical complete I-F system that consists of 6 genes (2). We then reconstructed a phylogenetic tree from all identified Cas7f protein sequences (Figure 2A, Supplemental Table S1, see respective Newick tree at ftp://ftp.ncbi.nih.gov/pub/makarova/supplement/Peters_at_al_2017/). Mapping gene neighborhoods on the tree revealed a single, monophyletic, strongly supported branch including all *cas1*-less I-F variants. As of this analysis, the branch encompassed 423 sequences from 19 genera of Gammaproteobacteria and appears to derive from a typical, complete I-F system (Figures 1A and 2A). Indeed, all other branches in the tree consist of Cas7f homologs from complete I-F systems containing a *cas1* gene within the locus. A few exceptions that are scattered in the tree are from either small contigs or disrupted *cas* loci. In the vast majority of the loci corresponding to the *cas1*-less branch, a *tniQ/tnsD* gene is located next to the *cas* genes (Figure 3).

**Figure 2.**
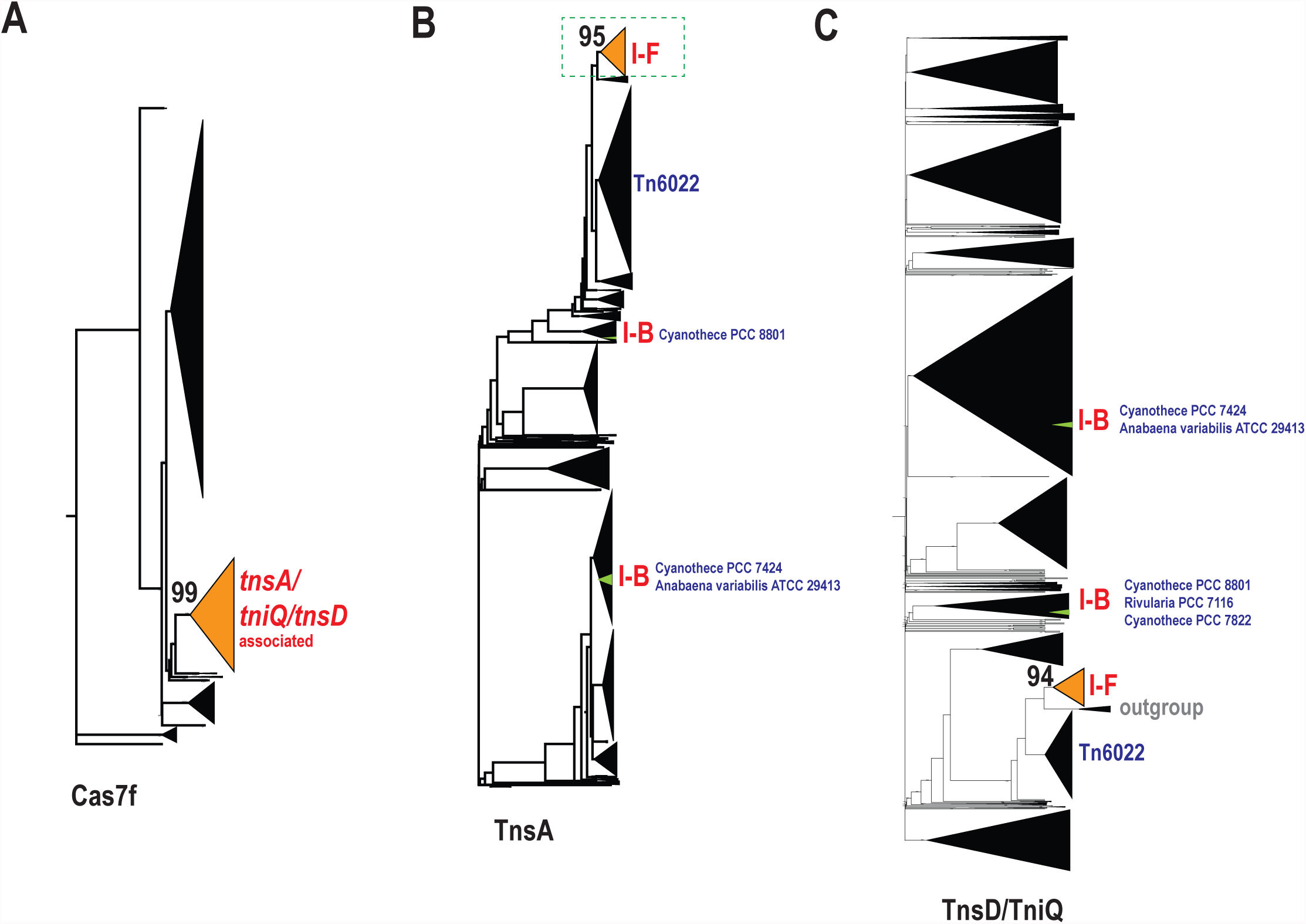
Schematic evolutionary trees for the Cas7f, TnsA and TniQ/TnsD protein families. **A.** The dendrogram was built using 2905 Cas7f proteins as described in Material and Method section (see complete tree at ftp://ftp.ncbi.nih.gov/pub/makarova/supplement/Peters_at_al_2017/). The major subtrees are collapsed and shown by triangles. The branch corresponding to the minimal I-F variant is colored in orange, and the bootstrap value for this subtree is shown. **B.** The dendrogram was built using 7023 TnsA protein sequences (see complete at ftp://ftp.ncbi.nih.gov/pub/makarova/supplement/Peters_at_al_2017/). The branch corresponding to TnsA in the loci containing I-F variant *cas* genes is colored in orange and I-B subtype *cas* genes are colored in green. The CRISPR-Cas subtypes are indicated next to the respective branches. Distinct cyanobacterial strains are indicated next to the respective I-B systems. The bootstrap value for TnsA branch associated with I-F *cas* genes is shown. **C.** The dendrogram was built using 7963 TniQ proteins (see complete tree at ftp://ftp.ncbi.nih.gov/pub/makarova/supplement/Peters_at_al_2017/). The designations are the same as for the TnsA dendrogram in B.

To determine whether the association of the Cas1-less I-F systems with Tn7-like elements was unique or emerged independently on several occasions, we analyzed the TniQ/TnsD and TnsA families. The TnsA protein is the most highly conserved gene of the Tn7-like elements and is responsible for the unique behavior of the elements with heteromeric transposases (31-34). We collected and annotated 10,349 loci containing at least *tniQ/tnsD* or *tnsA* (Supplemental Table S2) and reconstructed a tree for both protein families (Figure 2 B and C and see respective Newick trees at ftp://ftp.ncbi.nih.gov/pub/makarova/supplement/Peters_at_al_2017/). In both trees, the loci containing *cas* genes of the *cas1*-less I-F variant mapped to strongly supported monophyletic branches (Figure 2 B and C). Thus, phylogenetic analysis of both Cas7f and the associated transposon-encoded proteins reveals a strong link between a specific group of Tn7-like elements and a distinct variant of the subtype I-F CRISPR-Cas system. The Tn7-like elements in the clade that includes Tn6022 were identified as the outgroups to the respective branches in both the TnsA and TniQ/TnsD trees, suggesting that a member of the Tn6022 family is the ancestor of the CRISPR-associated variety of Tn7-like transposons (Figure 2B and C). Both clades include multiple, deep branches that are not associated with *cas* genes in the respective loci suggesting that the link with I-F system evolved relatively late in the history of this group of Tn7-like elements (see respective Newick trees at ftp://ftp.ncbi.nih.gov/pub/makarova/supplement/Peters_at_al_2017/). In several cases, however, the *cas* genes seem to have been lost from the vicinity of the conserved transposon genes (eg. *Shewanella baltica* OS678 and *Thiomicrospira crunogena* XCL_2), suggesting that the CRISPR-Cas system is not essential for the transposon survival. However, there are no intact *cas1*-less I-F systems outside this transposon neighborhood, with the implication that this CRISPR-Cas variant is functional only when associated with a Tn7-like element.

We further investigated the *tniQ/tnsD/tnsA* loci in order to identify any other CRISPR-Cas systems that might be linked to Tn7-like transposons. Only a few such instances were identified, mostly complete loci containing the adaptation genes. The respective *tnsA* and/or *tniQ/tnsD* genes are scattered in the phylogenetic trees suggesting that most of these associations are effectively random and might be transient (Supplemental Table S2). However, some of such loci do show a degree of evolutionary coherence. Specifically, they form two small, unrelated branches in both the TnsA and the TniQ/TnsD trees (See I-B in Figure 2B-C). All these CRISPR-*cas* loci are present in different cyanobacteria, belong to the I-B subtype and lack adaptation genes as well as the *cas3* gene that is required for DNA cleavage in type I systems. Thus, to a large extent, these type I-B variants mimic the organization of the more common transposon-associated, *cas1*-less I-F variant (See below).

### The *cas1*-less type I-F CRISPR-Cas system is mobilized together with conserved transposition genes

We analyzed the transposon end-sequences in the loci containing the I-F and I-B CRISPR-Cas variants in order to determine whether the *cas* genes were located within the boundaries of these elements or are simply adjacent to the transposon. The structure of the left and right ends of canonical Tn7 has been defined previously (Supplemental Figure S1). Tn7 ends are marked by a series of 22 bp TnsB-binding sites (35-37). Flanking the most distal TnsB-binding sites is an 8 base pair terminal sequence ending with 5′-TGT-3′/3′-ACA-5′. Tn7 contains 4 overlapping TnsB-binding sites in the ~90 bp right end of the element and three dispersed sites in the ~150 bp left end of the element, but the number and distribution of TnsB-binding sites can vary among Tn7-like elements (25, 31). End-sequences of Tn7-related elements can be determined by identifying the directly-repeated 5 base pair target site duplication, the terminal 8 base-pair sequence, and 22 base pair TnsB-binding sites (the latter two found in an inverted configuration in the left and right ends of the element) (Supplemental Figure S1). Compared with the canonical Tn7 and Tn6022, Tn7-like elements show extensive variation in size and gene complements as illustrated by a representative set of 12 complete elements ranging in size from 22 kb to almost 120 kb (38, 39)(Figure 3 and Table 1). One of these elements has been previously identified in *Vibrio parahaemolyticus* RIMD2210633 as a member of the Tn7 superfamily and encodes the *Vibrio* pathogenicity determinant, the thermostable direct hemolysin (TDH) (40).

**Figure 3.**
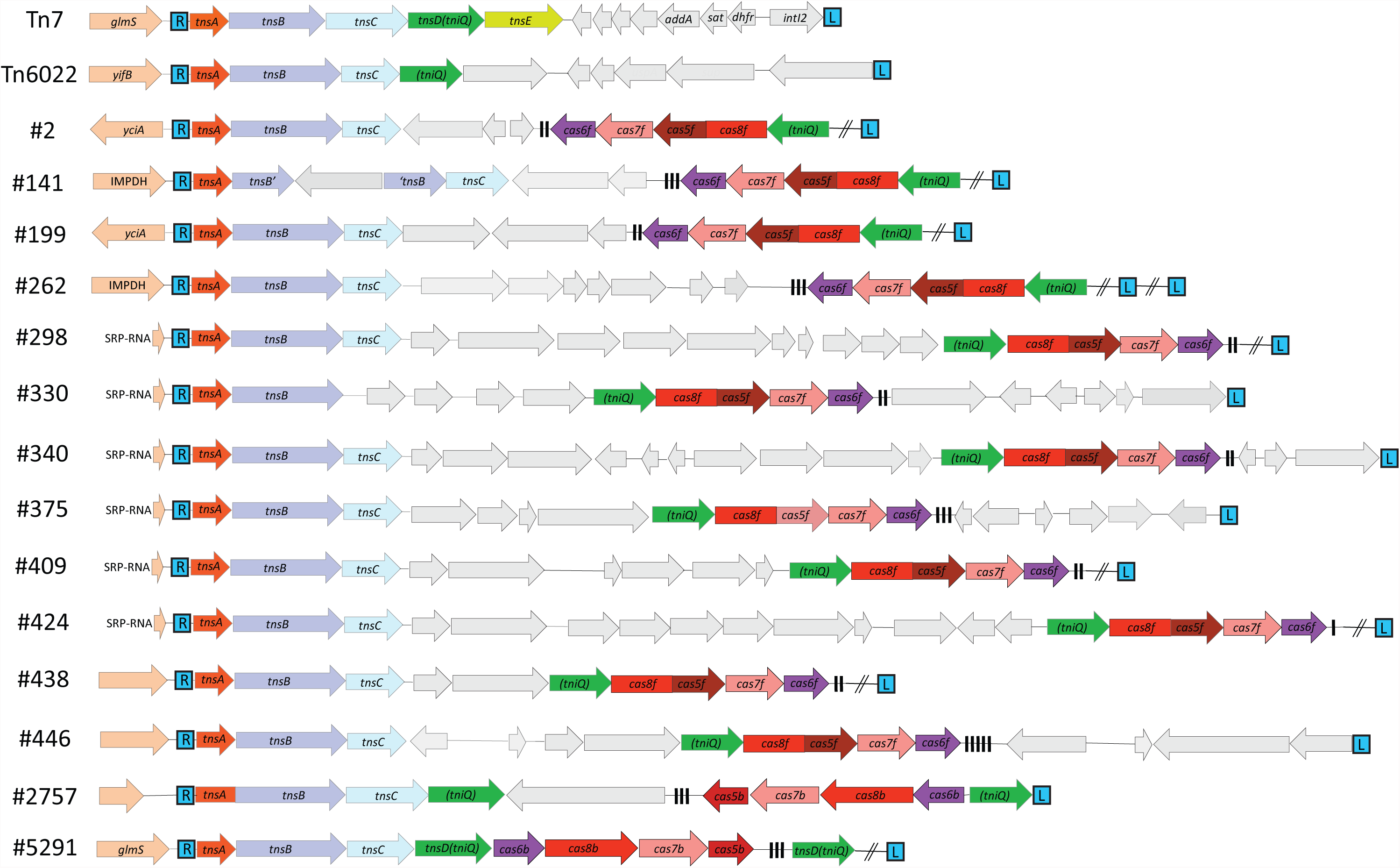
Schematic representation of Tn7, Tn6022, and selected Tn7-like transposons containing *cas* genes. Genomic features recognized by the transposon-encoded TniQ protein are indicated on the left (*glmS, yifB,* IMPDH, *yciA* and SRP-RNA). Color coding and labeling are as in Figure 1. Elements other than Tn7 and Tn6022 are denoted by the respective TnsA tree leaves (#XX)(Tn6022=Tree node #582)(Supplemental Table S2). Other genes are shown in grey, and known Tn7 cargo genes are indicated. Black vertical bars indicate repeats in the element-encoded arrays. DNA sequences omitted in the graphic is shown with two back slashes. See text for details.

**Table 1.**
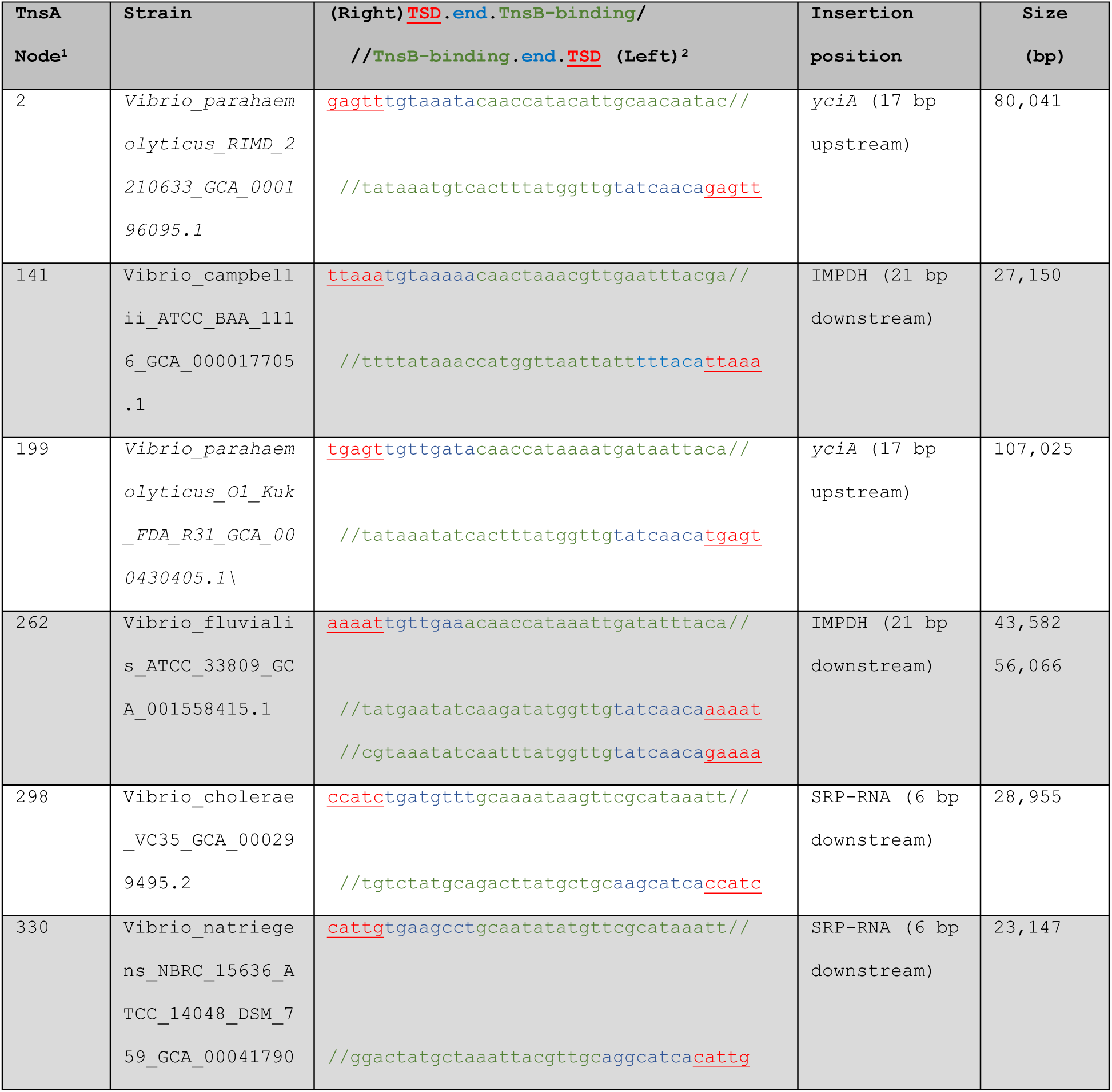
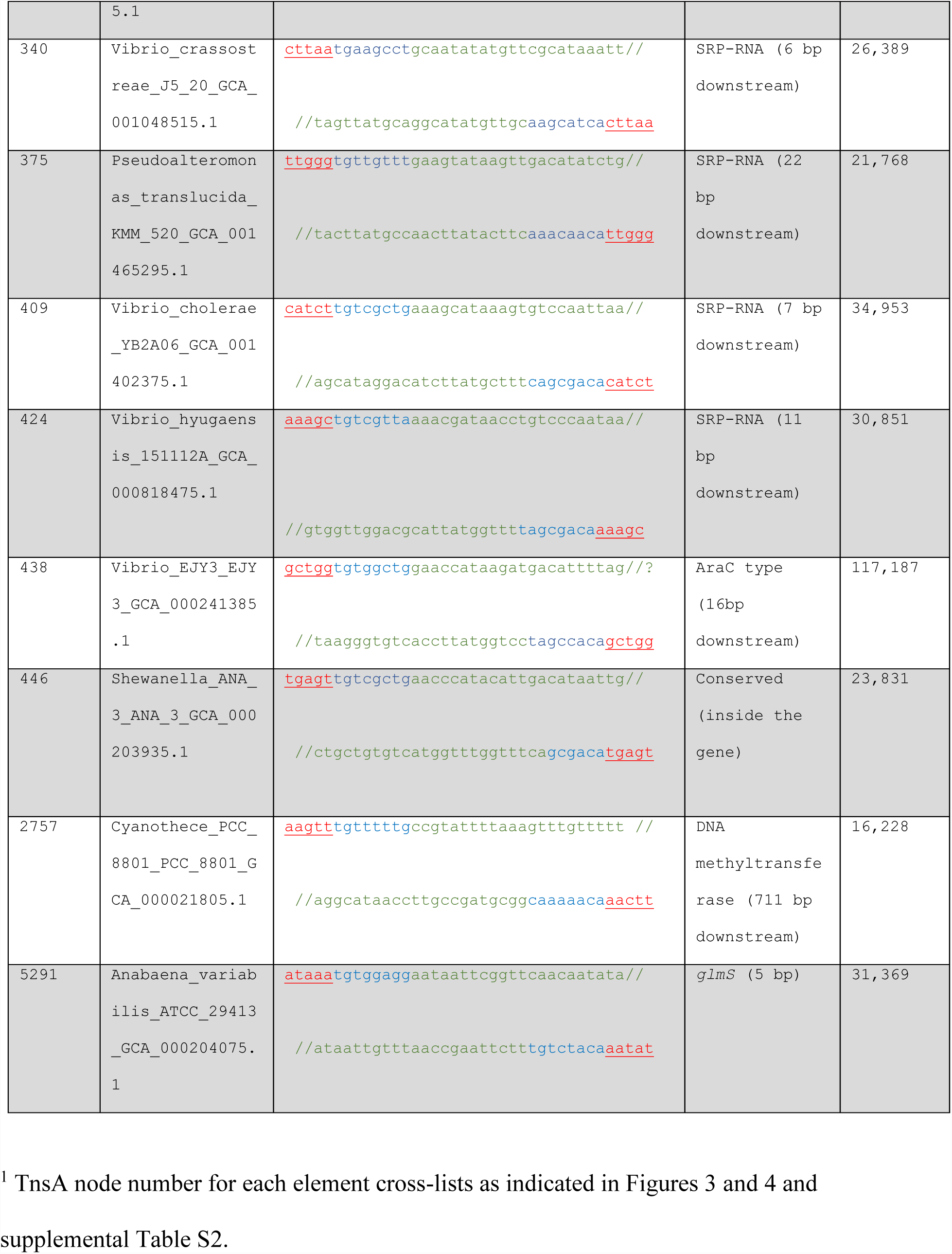
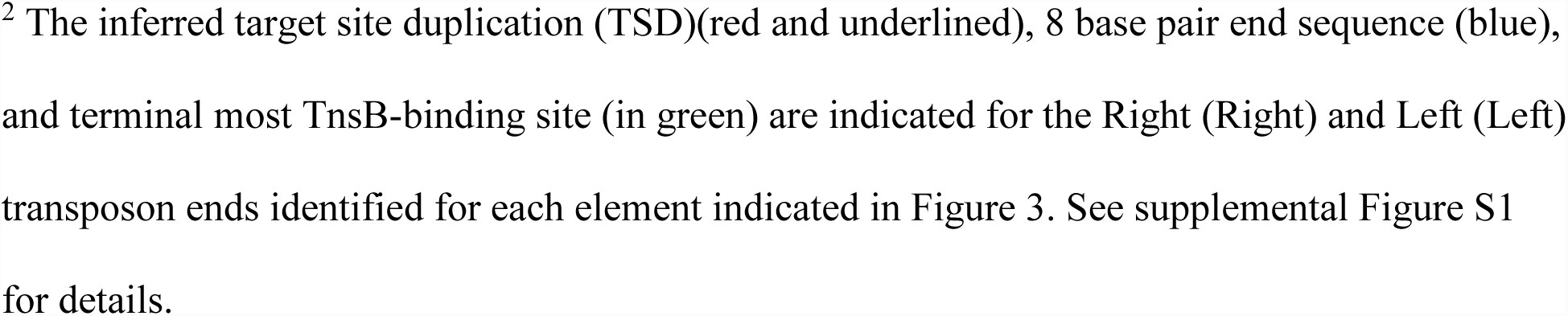
Characteristics of selected Tn7 loci associated with *cas* genes.

In our analysis of CRISPR-Cas systems, two groups of type I-B variants were identified in association with Tn7-like elements (Figure 2B-C). Similar to the type I-F CRISPR-Cas variant, these I-B systems are expected to be functional for maturing CRISPR transcripts and forming crRNA complexes at protospacers but lack adaptation genes and Cas3, and accordingly, are likely to be defective for interference. Furthermore, these type I-B CRISPR-Cas variants are associated with small CRISPR arrays (Figure 3). One group of the type I-B associated transposons encodes a TnsD/TniQ protein that belongs to the same clade as TnsD from canonical Tn7 and resides in the *attTn7* site downstream of *glmS*. An example from this subgroup has been previously identified in *Anabaena variabilis* (# 5291 in Figure 3), but the minimal Cas system contained in the element was not analyzed (41). The second group encodes a TniQ protein that belongs to a new family of elements encoding fused TnsA-TnsB proteins (#2757 in Figure 3).

Taken together, these findings indicate that the type I-F and I-B CRISPR-Cas variants identified in this work are part of the core gene repertoire in multiple clades of Tn7-like elements.

### Chromosomal insertions of the I-F CRISPR-Cas-associated elements show three recognizable attachment sites likely accessed by dedicated TniQ/TnsD proteins

The canonical Tn7 element has been studied extensively, especially the transposition pathway that directs the element into the *attTn7* site located downstream of the conserved *glmS* gene. The Tn7 TnsD(TniQ) protein is a sequence-specific DNA-binding protein that recognizes a highly conserved 36 bp sequence in the downstream region of the *glmS* gene coding sequence (26, 42). Transposition events promoted by TnsABC+D are directed into a position 23 bp downstream of the region bound by TnsD. Tn7 transposition is orientation-specific in all transposition pathways; the transposon end proximal to the *tnsA* gene (the “right” end of the element) is adjacent to the DNA sequence or a specific protein complex recognized in each pathway (29, 42-44).

We analyzed the region adjacent to the point of insertion of the elements and identified three new attachment sites for the *cas1*-less, type I-F-associated transposons. Similar to Tn7 insertions, one subgroup of the elements occurred downstream of a gene, but instead of *glmS*, these insertions were found downstream of an inosine-5′-monophosphate dehydrogenase gene (Table 1, Figure 3 and 4). The configurations found with the other recognizable attachment sites were new for Tn7-like elements. In one case, the attachment site was located upstream of the *yciA* gene, which encodes an acyl-CoA thioester hydrolase (Table 1, Figure 3 and 4). Presumably, insertion of the element into this attachment site would ablate the normal promoter, but changes in expression remain to be demonstrated experimentally. The third attachment site identified for the *cas1*-less type I-F-associated elements is the first example where a non-proteinen-coding gene was recognized, namely, the gene for the signal recognition particle RNA (SNP-RNA) (Table 1, Figure 3 and 4). The concordance between the phylogeny of the TniQ/TnsD proteins and the attachment site used by the element is consistent with the hypothesis that each attachment site is recognized by a cognate TniQ/TnsD protein (Figure 4). Despite this concordance, many transposons appear to be inserted in random sites (Figure 4). It remains unclear how insertions were directed into these sites because they are unlikely to be specifically recognized by TniQ/TnsD proteins encoded by these elements, and these elements lack a homolog of the TnsE protein found in typical Tn7 transposons.

**Figure 4.**
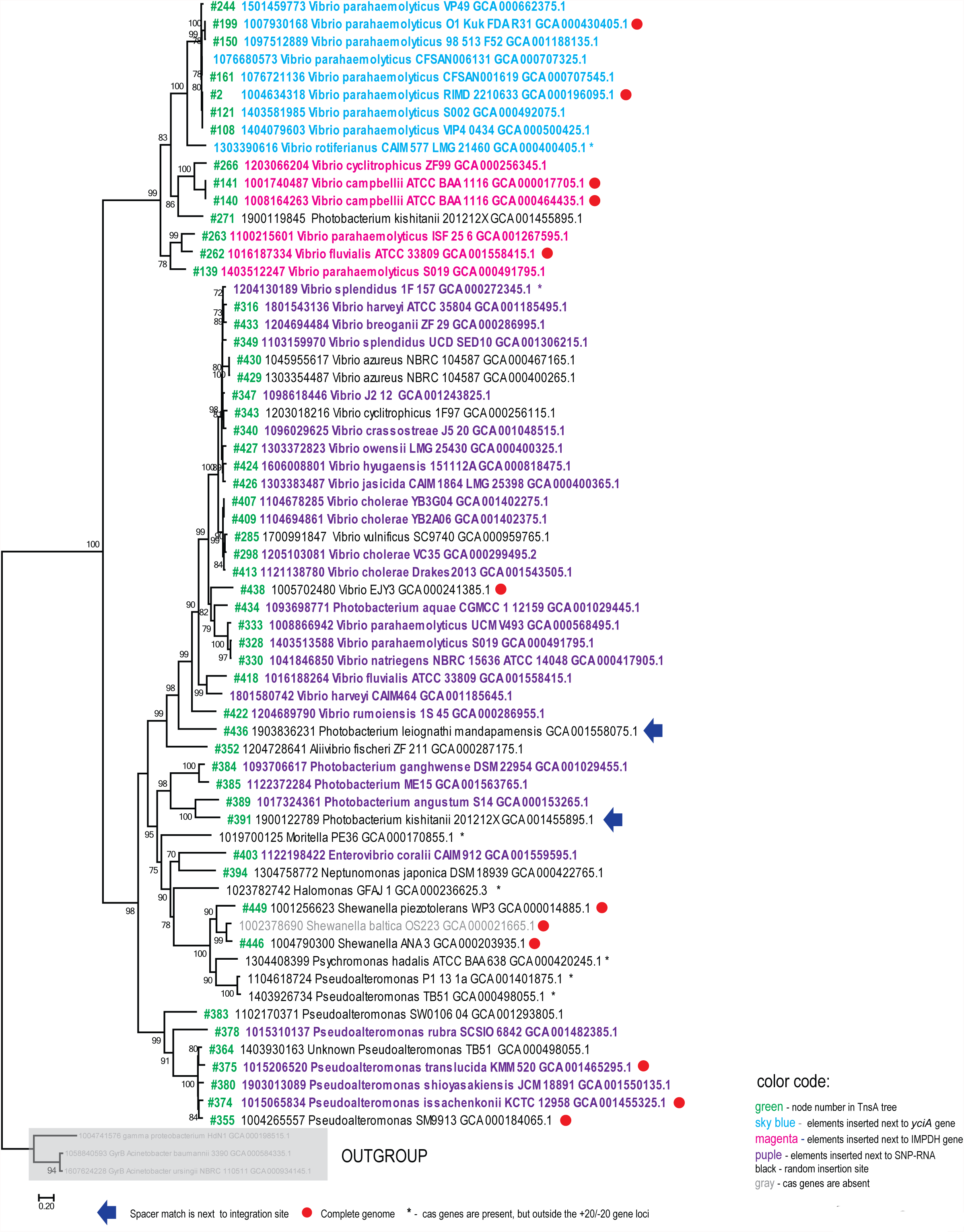
Phylogenetic tree of selected representatives of type I-F-associated TniQ-like proteins. A maximum likelihood phylogenetic tree was built as described in Materials and Methods for a selected set of TniQ-like proteins associated with the type I-F CRISPR-Cas variant and a few sequences from the outgroup outlined in the Figure 2C (67 sequences altogether). The numbers at internal branches indicate bootstrap support (percent); only values greater than 70% are indicated. Elements located in one of the three attachment sites identified in this work are shown by color as indicated (*yciA*, IMPDH, and SRP-RNA); random sites. The TniQ tree leaves (#XX)(Supplemental Table S2) are shown in green.

### Analysis of CRISPR arrays associated with the *cas1*-less I-F systems

The great majority of the transposon-associated I-F and I-B systems encompass a CRISPR array downstream of the *cas6* gene (See Figure 3 for examples). In most cases, this array contains only one or two spacers, suggesting that spacer acquisition in these arrays occurs only rarely (Table 2 and Figure 3). Nevertheless, the spacers are typically unrelated even in closely related bacterial genomes indicating that, occasionally, new spacers are incorporated, and old ones are lost. Obviously, only adaptation genes acting *in trans* can insert new spacers into these arrays. Among the 14 complete bacterial genomes containing elements with the I-F CRISPR-Cas, only two encompass other CRISPR-Cas loci containing adaptation genes, namely, *Vibrio fluvialis* ATCC 33809 and *Pseudoalteromonas rubra* SCSIO6842 that possess I-F and I-C systems, respectively. Among draft genomes, there are more cases where additional, complete CRISPR-Cas systems, mostly I-F and I-E, are present in the same genomes. Nevertheless, most of the genomes that contain the I-F variant associated with Tn7-like transposons lack other CRISPR-Cas system that would be able to provide for adaptation, which might account for the short CRISPR arrays. All 4 complete genomes containing elements associated with I-B systems encompass additional CRISPR-Cas loci containing adaptation genes, often of subtype I-D, which is abundant in cyanobacteria (2).

**Table 2.**
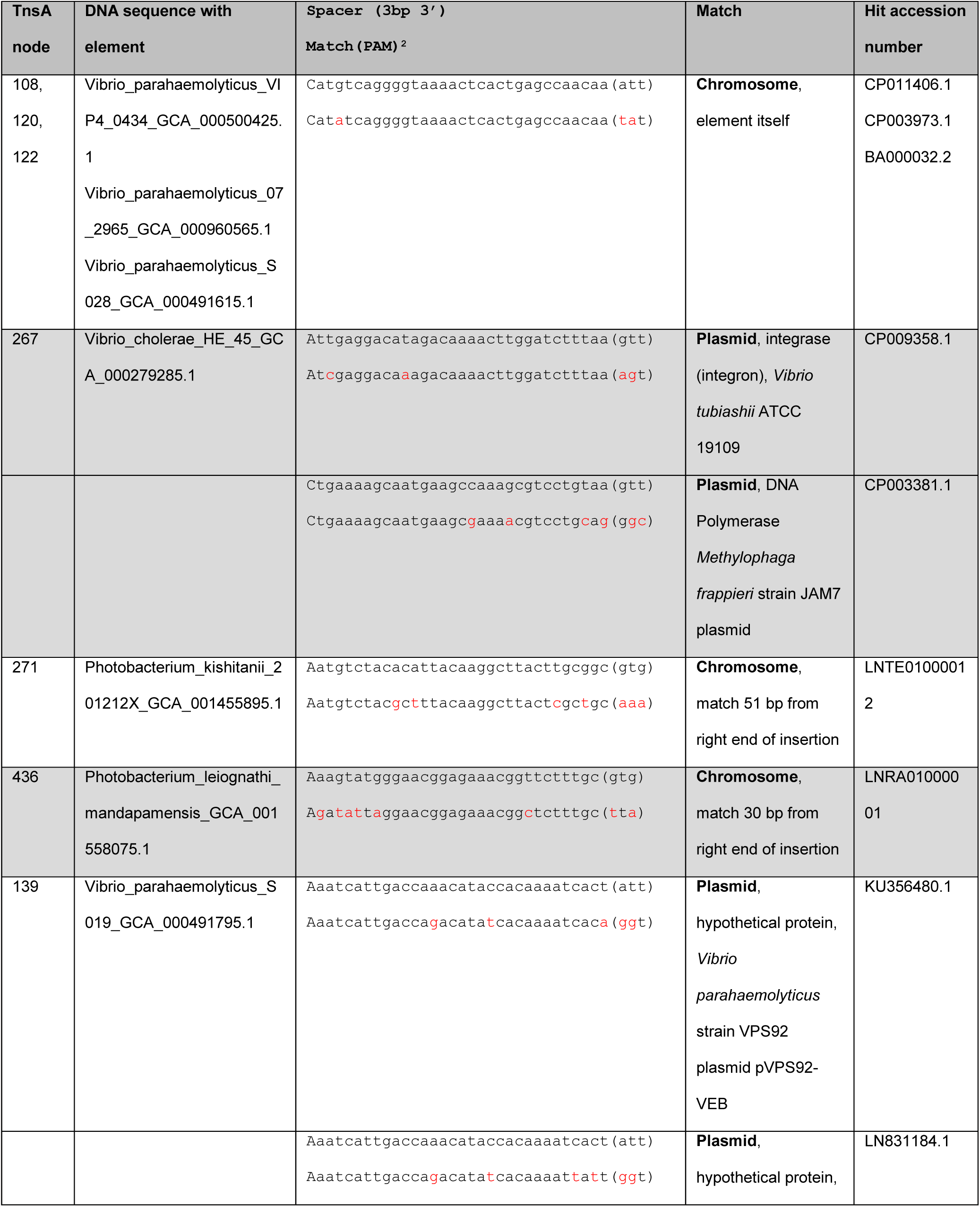
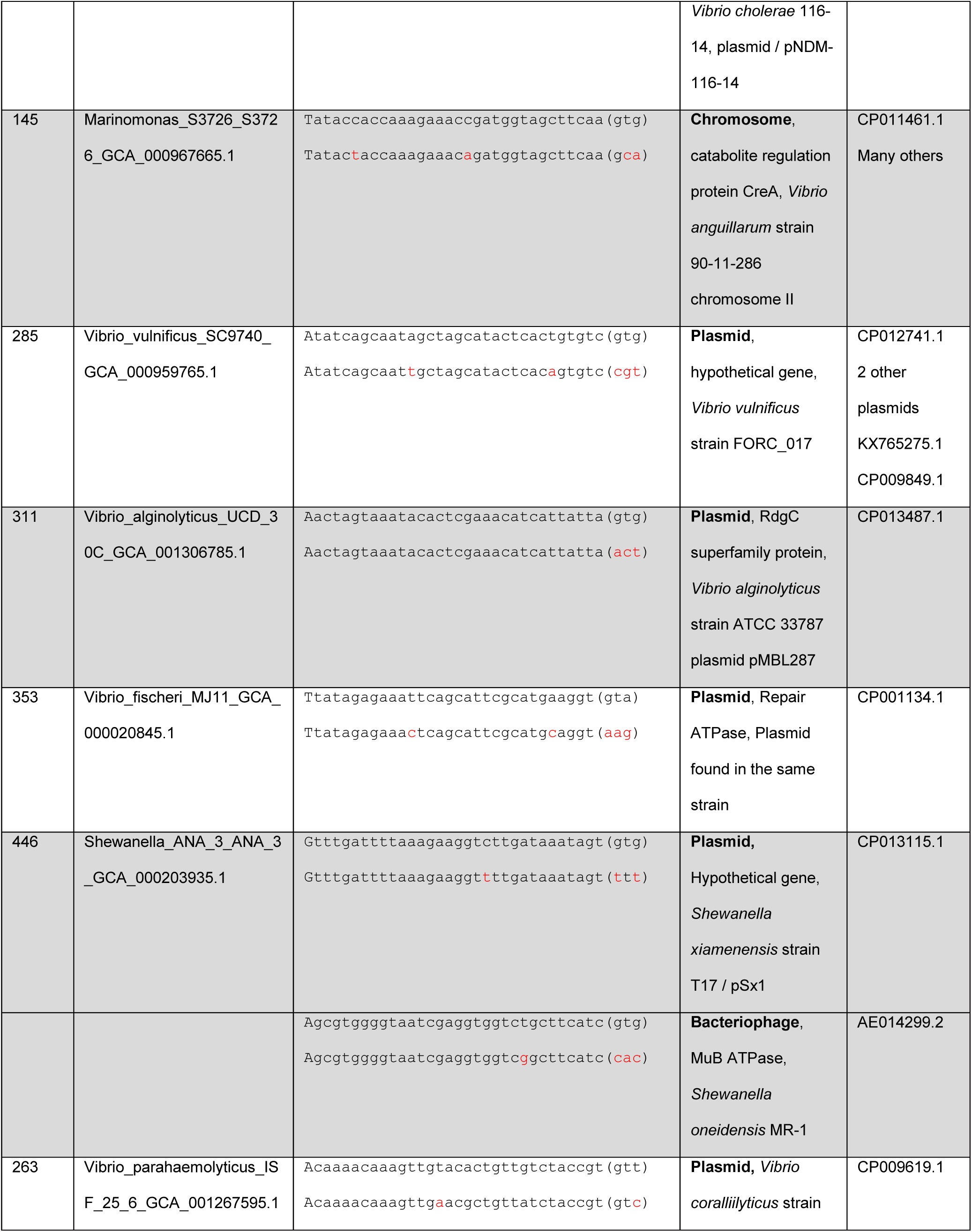
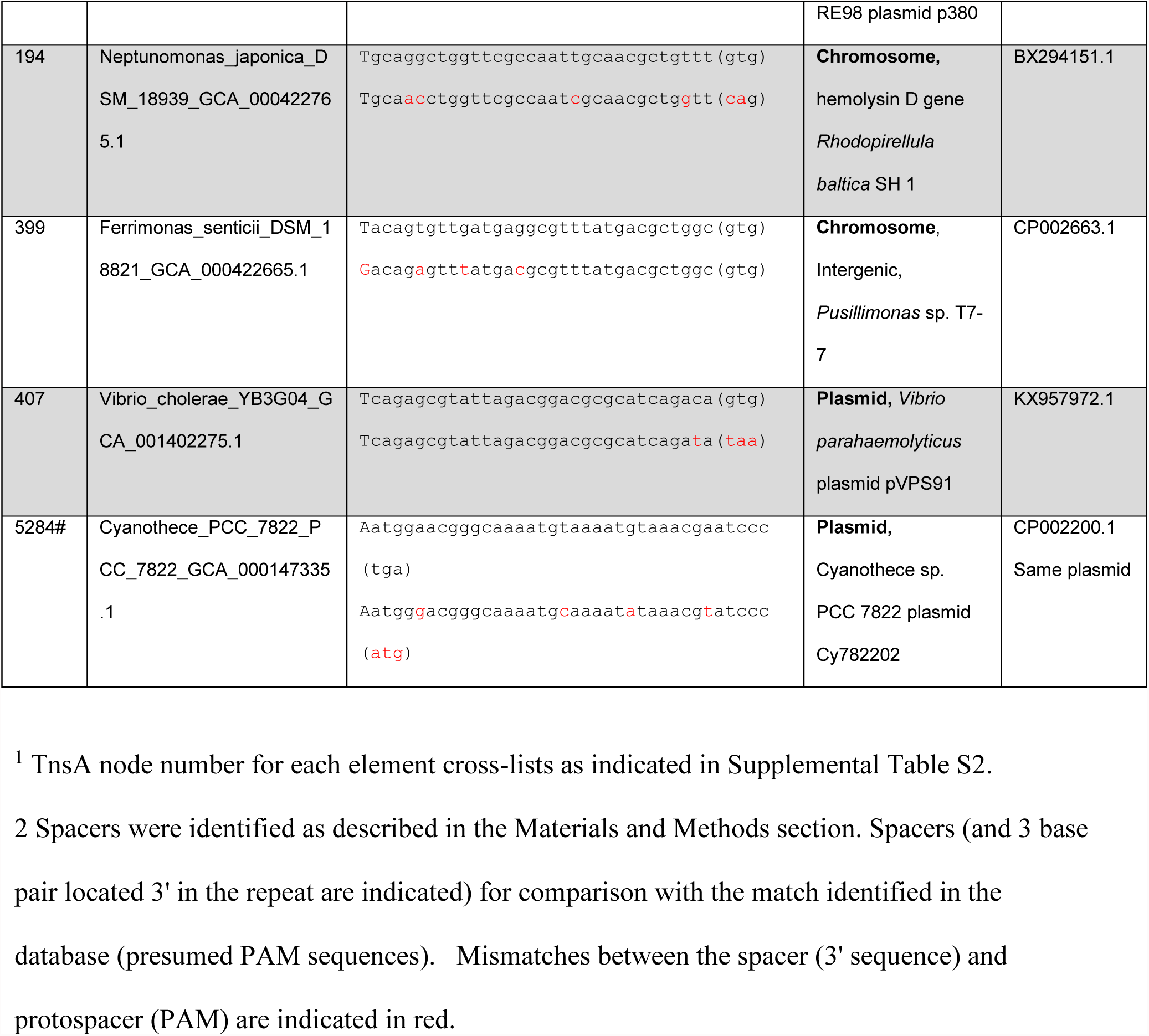
Spacers with identified matches.

Altogether, more than 800 spacers were identified in the transposon-associated I-F and I-B CRISPR arrays (see automatically and manually identified spacers at ftp://ftp.ncbi.nih.gov/pub/makarova/supplement/Peters_at_al_2017/). As in most analyses of CRISPR spacers (45-47), only a small fraction of these spacers yielded significant matches to sequences in public databases. However, the matches that could be detected were informative because they were to plasmids and bacteriophages associated with the same bacterial genera where the respective elements are found (Table 2). We identified two cases (in *Photobacterium kishitanii* and *Photobacterium leiognathi*) of potential special interest, where spacers matched the region adjacent to the *tnsA*-gene proximal side of the element (Table 2), i.e. the specific region where complexes involved in targeting transposition events interact with the target DNA (29, 48, 49). An additional spacer match was found inside the transposon boundaries in several *Vibrio parahaemolyticus* strains (Table 2). A similar situation might have also occurred in a Tn7-like transposon associated with a type I-B CRISPR-Cas variant in a *Cyanothece* PCC 7822 plasmid although end sequences could not be unambiguously defined for this element (Table 2).

### A role for CRISPR-Cas in targeting transposition?

Taking into account all the observations on the transposon-associated CRISPR-Cas systems and previous studies on the mechanism of target site activation, we propose a model for the involvement of Cas1-less CRISPR-Cas systems in targeting transposition to facilitate cell-cell transfer of the element (Figure 5). Canonical Tn7 encodes two targeting pathways that are both mediated by the same set of TnsABC proteins (Figure 1B). The TnsABC+TnsD(TniQ) pathway appears to be broadly conserved allowing high frequency transposition into an attachment site recognized by a cognate TnsD/TniQ protein (Figure 1 and 4, Table 1)(31). The *cas1*-less I-F CRISPR-Cas variant is encoded in the same location where the *tnsE* gene that promotes transposition into conjugal plasmids and filamentous bacteriophages is typically located (Figure 3). Thus, it appears plausible that the CRISPR-Cas system functionally replaces *tnsE* as a mechanism facilitating horizontal transfer of the element. Support for this possibility comes from the observation that the transposon-associated CRISPR arrays largely carry plasmid and phage-specific spacers and could direct the transposon to the respective elements (Table 2).

**Figure 5.**
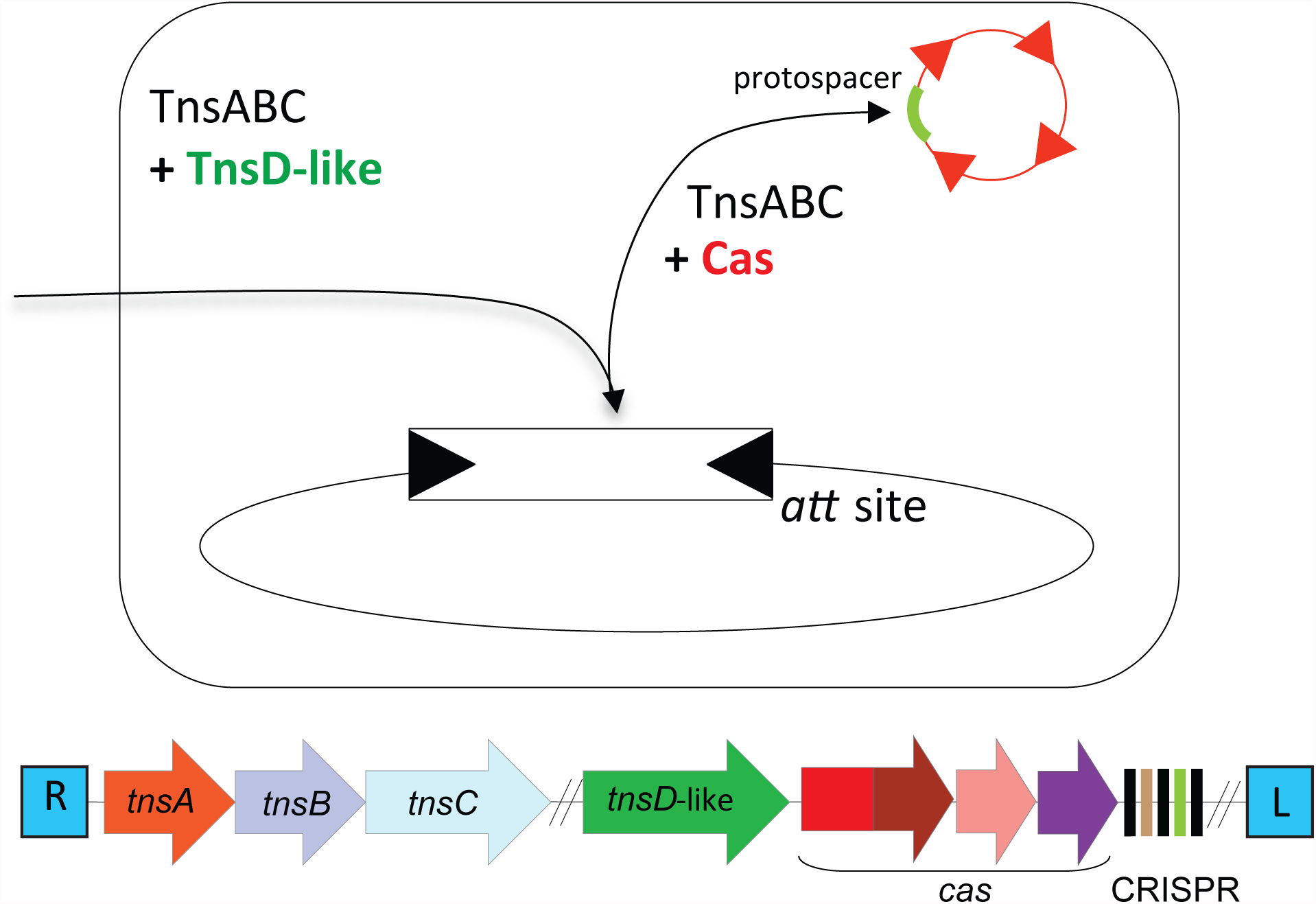
Model of the two targeting pathways for Tn7 elements containing CRISPR-Cas system. Designations are the same as in Figure 1.

### Distortions in B form DNA induced by Cas-crRNA could play a role in recruiting transposition

Transposition into *attTn7* is well-understood at the molecular level; the DNA structure in the vicinity of the attachment site plays a central role in transposition (Figure 6 A, B and C). TnsD-binding induces an asymmetric distortion in the *attTn7* target DNA that is primarily responsible for attracting TnsC during transposition (48, 50)(Figure 6A). Protein-protein interaction between TnsD and TnsC have been detected (26) but multiple lines of evidence suggest a defining role for the distortion in the DNA for the target site selection (see below).

**Figure 6.**
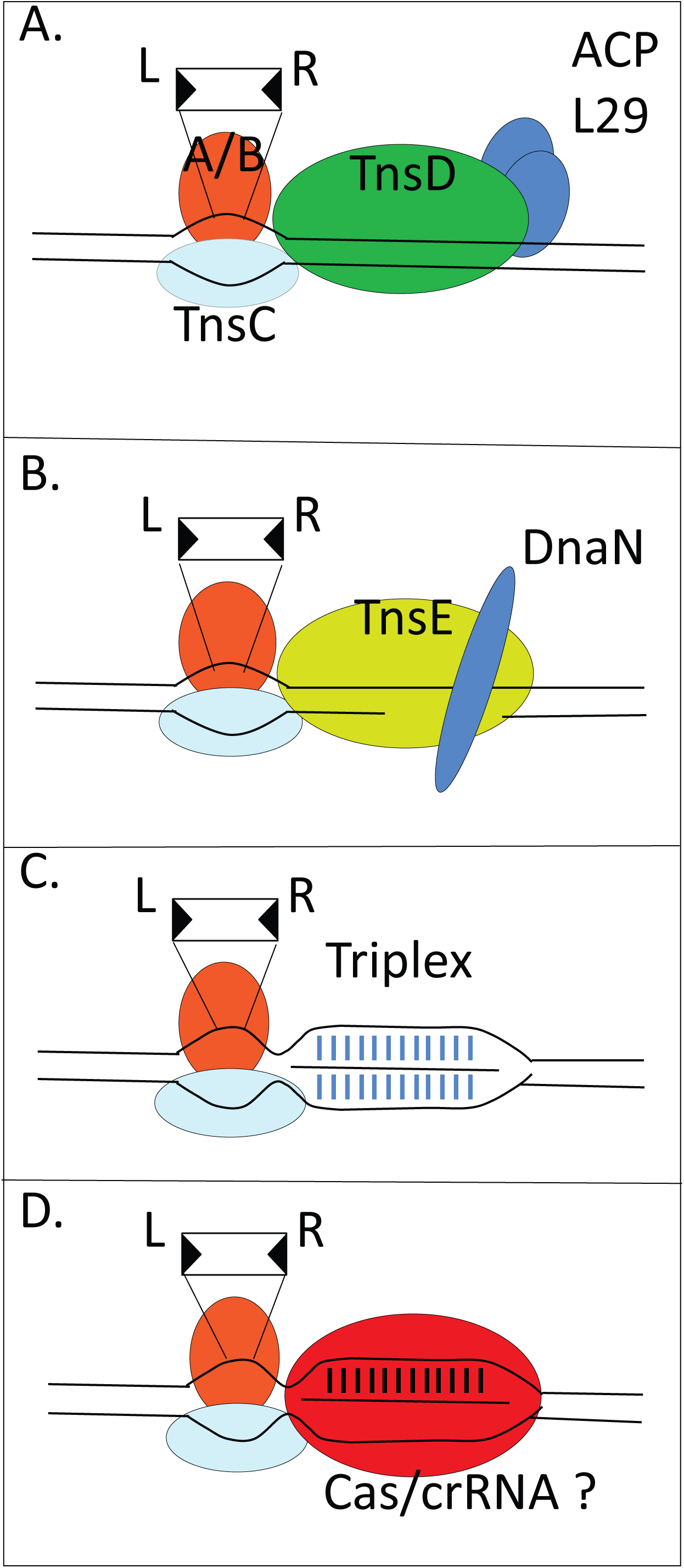
Models of the three previously described Tn7 targeting pathways and the proposed CRISPR-Cas-facilitated transposition pathway. Representations of TnsABC+TnsD (A) and TnsABC+TnsE (B) transposition pathways, the synthetic transposition pathway that targets triplex DNA complexes with a mutant form of TnsC, TnsABC* (C) and the proposed targeting pathway mediated by Cas interference complexes (D) are shown. Known host factors that participate in the TnsD and TnsE pathways are also shown (ACP, L29, and DnaN). See text for details and references.

The TnsABC proteins are normally insufficient for Tn7 transposition *in vivo* or *in vitro* (51); however, certain gain-of-activity mutations in the regulator protein TnsC (TnsC*) allow transposition in the absence of TnsD or TnsE (52, 53). Untargeted transposition is observed *in vitro* and *in vivo* with TnsABC* in the absence of target site selection proteins (30, 52), but notably, transposition in this case is attracted to a specific location adjacent to a short segment of triplex-forming DNAs (44, 54). Analogous to transposition events found in *attTn7*, these events are targeted to a position on one side of the triplex-forming DNA in a unique orientation owing to the ability of TnsC to recognize the distortion formed at the triplex-to-B-form DNA transition (Figure 6C). Distortions induced in the target DNA are also implicated in transposition targeting by TnsABC+E (55) (Figure 6B). Given that distortions in B form DNA are also expected adjacent to crRNA-bound effector complexes that generate R-loops through duplex formation between the crRNA and the protospacer (12, 56), there could be a mechanistic link between the well-understood Tn7 targeting process and DNA targeting by the CRISPR-Cas effector complexes (Figure 6D). This analogy is consistent with the left to right orientation of the two insertions located adjacent to spacer matches (Table 2).

### Evolution of the association between CRISPR-Cas variants and Tn7 like elements

Given that at least some type I CRISPR-Cas systems selectively integrate spacers from plasmids and phages (6, 57), an attractive hypothesis is that the CRISPR-*cas* loci that randomly became associated with the transposon were fixed through selection for their ability to facilitate dissemination of transposons. As discussed above, because changes in DNA structure play an important role in target site selection by Tn7, relatively little evolutionary adaptation might be needed to allow the core TnsABC machinery to recognize crRNA-bound effector complexes for targeting. In this light, it is intriguing that association between CRISPR-Cas systems and Tn7-like elements occurred on multiple, independent occasions. The consistent minimalist features in the organization of the transposon-associated type I-F and I-B variants imply that they co-evolved with Tn7-like elements along parallel paths of reductive evolution. Both type I-F and type I-B systems have lost the adaptation module (*cas1* and *cas2*) and the *cas3* gene, which is required to cleave the target DNA in other type I systems (1). The absence of Cas3 implies that these CRISPR-Cas systems recognize but do not cleave the target, a mode of action that would allow the targeted DNA to serve as a vehicle for horizontal transfer of the respective Tn7-like transposon.

The transposon-associated CRISPR arrays are short, and the respective bacterial genomes often lack CRISPR adaptation modules. Thus, the majority of the CRISPR-containing transposons are likely to be relatively recent arrivals to the respective genomes, conceivably, brought about by the plasmid or phage against which they carry a spacer. Once integrated into a new host attachment site, such transposons could “lie in waiting” for a horizontal transfer vehicle, either as a result of *in trans* acquisition of a new spacer that is specific to an endogenous plasmid or prophage or via the entry of an element that is already represented by a cognate spacer in the transposon-encoded CRISPR array. In some cases, an incoming plasmid or phage recognized by the CRISPR-Cas system and targeted for transposition would be incapacitated by the integration event. Nevertheless, even such unproductive integrations would still benefit the CRISPR-carrying transposon by protecting the host. In such cases, CRISPR-directed integration that is in keeping with a selfish behavior for the transposon would also qualify as altruistic behavior toward the host. It is unclear what advantage there could be to losing the capacity to undergo adaptation (i.e. lack of *cas1* and *cas2*). However, because Tn7 itself is a mobile element, sporadic access to adaptation systems *in trans* may prove safer for the element, limiting the acquisition of transposon-directed spacers in the element-encoded array. Occasionally, these elements appear to acquire spacers from the host chromosome, conceivably stimulating ectopic transposition within the same genome. Such a system could be beneficial in allowing transposition in hosts that lack attachment sites recognized by the element-encoded TniQ/TnsD protein.

Many questions remain regarding the functioning of the CRISPR-Cas in Tn7-like transposons including the possibility of direct interaction between the CRISPR effector complexes and either TnsD/TniQ, TnsABC, or other transposon-encoded accessory proteins. It is also unclear if these CRISPR-Cas variants might perform alternative or additional functions, beyond facilitation of transposition, such as gene silencing or protection of the transposon.

From the evolutionary standpoint, the transposon-associated CRISPR-Cas systems fit the “guns for hire” paradigm (58). Under this concept, genes of mobile genetic elements (MGE) genes are often recruited by host defense systems, and conversely, defense systems or components thereof can be captured by MGE and repurposed for counter-defense or other roles in the life cycle of the element. Recruitment of MGE apparently was central to the evolution of CRISPR-Cas, contributing to the origin of both the adaptation module and the Class 2 effector modules (3, 56, 59). On the other side of the equation, virus-encoded CRISPR-Cas systems have been identified and implicated in inhibition of host defense (60). The observations described here, if validated experimentally, seem to “close the circle” by demonstrating recruitment of CRISPR-Cas systems by transposons, conceivably, for a role in targeting transposition, a key step in transposon propagation. Finally, it has not escaped our notice that the transposon-encoded CRISPR-Cas systems described here potentially could be harnessed for genome engineering applications, namely, precise targeting of synthetic transposons encoding selectable markers and other genes of interest.

## Methods

### Prokaryotic Genome Database and open reading frame annotation

Archaeal and bacterial complete and draft genome sequences were downloaded from the NCBI FTP site (ftp://ftp.ncbi.nlm.nih.gov/genomes/all/) in March 2016. For incompletely annotated genomes (coding density less than 0.6 CDS per kbp), the existing annotation was discarded and replaced with the Meta-GeneMark 1 (61) annotation using the standard model MetaGeneMark_v1.mod (Heuristic model for genetic code 11 and GC 30). Altogether, the database includes 4,961 completely sequenced and assembled genomes and 43,599 partially sequenced genomes.

Profiles for three families protein families, namely Cas7f (cd09737, pfam09615), TnsA (pfam08722, pfam08721) and TnsQ/TnsD (pfam06527), that are available in the NCBI CDD database (62) were used as queries for PSI-BLAST searches (E-value: 10^−4^, other parameters were default) to find respective homologs. All ORFs within 10 kb regions up- and downstream of *cas7f* genes (to cover potential complete I-F system) and 20 kb regions up- and downstream of *tnsQ/tnsD* and *tnsA* (to cover potential Tn7-like elements) were further annotated using RPS-BLAST searches with 30,953 profiles (COG, pfam, cd) from the NCBI CDD database and 217 custom Cas protein profiles (2). The CRISPR-Cas system (sub)type identification for all loci was performed using previously described procedures (2).

### Protospacer analysis

The CRISPRfinder (63) and PILER-CR (64) programs were used with default parameters to identify CRISPR arrays in Cas7f and TnsA/TnsD loci. The MEGABLAST program (65) (word size 18, otherwise default parameters) was used to search for protospacers in the virus subset of the NR database and the prokaryotic genome database. Matches were considered only if they showed at least 95% identity and at least 95% length coverage in the case of the NR database, and 80% identity and 80% length coverage for the self-hits (hits were classified as self if they matched the same genomes or genome of the same species disregarding the strain information). Because the automatic approach missed several short CRISPR arrays, loci initially found to lack CRISPR were analyzed manually by examining the intergenic region downstream of the *cas6f* gene for repeats and using the BLASTN program with the default parameters to find matches to the spacer identified.

### Clustering and Phylogenetic Analysis

To construct a non-redundant, representative sequence set, protein sequences within families of interest were clustered using the NCBI BLASTCLUST program. (ftp://ftp.ncbi.nih.gov/blast/documents/blastclust.html) with the sequence identity threshold of 90% and length coverage threshold of 0.9. Short fragments or disrupted sequences were discarded. Multiple alignments of protein sequences were constructed using MUSCLE (66) or MAFFT (67) programs. Sites with the gap character fraction values >0.5 and homogeneity <0.1 were removed from the alignment. Phylogenetic analysis was performed using the FastTree program (68), with the WAG evolutionary model and the discrete gamma model with 20 rate categories. The same program was used to compute bootstrap values.

Relationships within diverse sequence families were established using the following procedure: initial sequence clusters were obtained using UCLUST (69) with the sequence similarity threshold of 0.5; sequences were aligned within clusters using MUSCLE (66). Then, cluster-to-cluster similarity scores were obtained using HHsearch (70) (including trivial clusters consisting of a single sequence each), and a UPGMA dendrogram was constructed from the pairwise similarity scores. Highly similar clusters (pairwise score to self-score ratio >0.1) were aligned to each other using HHALIGN (70), and the procedure was repeated iteratively. At the last step, sequence-based trees were reconstructed from the cluster alignments using the FastTree program (68) as described above and rooted by mid-point; these trees were grafted onto the tips of the profile similarity-based UPGMA dendrogram.

### Analysis of Tn7-like elements

End-sequences of Tn7-like elements were determined by identifying the directly-repeated five base pair target site duplication, the terminal eight base-pair sequence, and 22 base pair TnsB-binding sites as described in the text using Gene Construction Kit 4.0 to manipulate DNA sequences and search for DNA repeats. Sequence files were derived from matches to *cas7f, tnsA* and *tniQ* as described above.

## Author contributions

J.E.P, K.S.M. and S.S. performed genomic analysis. J.E.P, K.S.M and E.V.K. designed the analysis, participated in the data interpretation and discussion, and wrote the paper.

## Acknowledgements

JEP was supported by the USDA National Institute of Food and Agriculture, Hatch project NYC-189438. KSM, SS and EVK are supported by the intramural program of the U.S. Department of Health and Human Services (to the National Library of Medicine).

## Competing interests

The authors declare no competing financial interests.

---

**Supplemental Figure S1.**
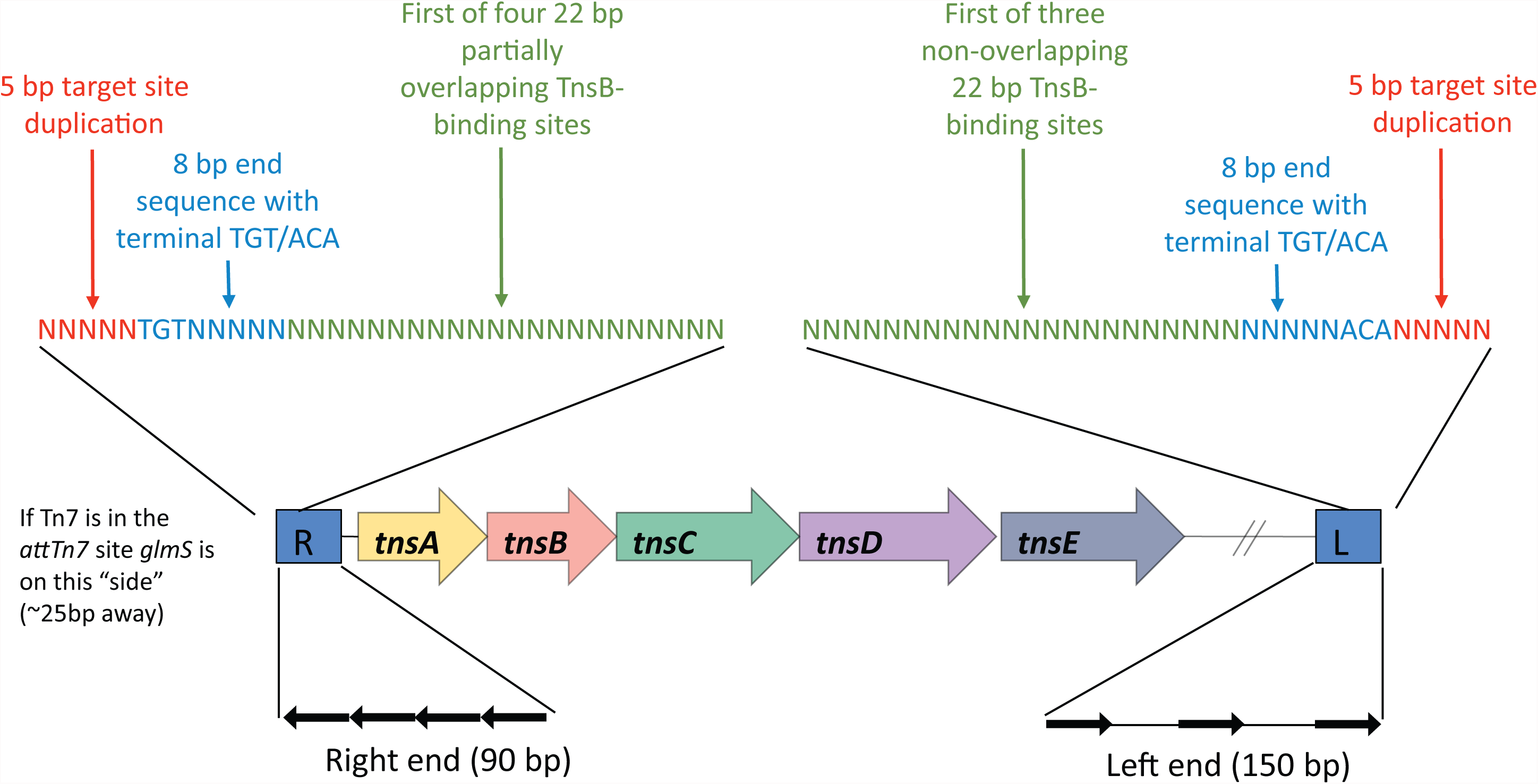
Schematic representation of the end structure of Tn7-like elements – Anatomy of a Tn7 insertion. Typically the insertion occurs at a single site about 25 bp downstream from the last codon of glmS. The Tn7 end proximal to tnsA is closest to glmS (by convention it is referred to as the “right” end). Transposition generates a target site duplication (shown in red) of the chromosomal sequence that now forms a direct repeat on either side the element. In the case of insertion of the canonical Tn7element at the attTn7 site this sequence is of GCGGG. There will be an 8 bp “end sequence” that starts with TGT/CAC, immediately after this will be the first 22 bp binding site for TnsB.

